# Nucleosome Core Allostery Governs Chromatin Recognition and Cell Fate

**DOI:** 10.64898/2026.09.24.753912

**Authors:** AN Wilkening, M Priyadarshini, MJ Trnka, MMK Wong, A Nawar, J Medwid, DN Kahan, S Marqusee, S Sanulli

## Abstract

Nucleosomes regulate chromatin folding, accessibility, and factor recruitment. Current models primarily attribute these functions to histone tail modifications, while the core is largely viewed as a structural scaffold. Yet subtle changes within the nucleosome core can produce profound functional consequences, and the mechanisms underlying these effects remain unclear. Here, we describe nucleosome core allostery as a fundamental principle of chromatin regulation that amplifies the impact of minimal nucleosome variations. Leveraging natural differences between H2A.Z variants, we show that the nucleosome core encodes distinct conformational dynamics that propagate allosterically, thereby controlling nucleosome accessibility and recognition by chromatin factors. As a result, a single buried amino acid substitution alone is sufficient to reprogram nucleosome dynamics and bias cell identity. Our findings establish the nucleosome core as an allosteric regulatory module and provide a generalizable framework for how subtle variation within nucleosomes is amplified into diverse biological outcomes in development and disease.

## Introduction

Nucleosomes, composed of ∼147 base pairs of DNA wrapped around a histone octamer, are the fundamental units of chromatin^1^. Beyond packaging the genome, nucleosomes function as regulatory modules that control DNA accessibility and the recruitment of chromatin-associated factors. Chromatin regulation has traditionally been attributed to modifications of the flexible histone tails, which modulate chromatin architecture and serve as recruitment platforms^2^. However, across diverse biological contexts, subtle differences in the nucleosome core are associated with profound cellular and organismal phenotypes, suggesting that the globular nucleosome core can also encode regulatory effects^3–7^.

Single amino acid changes in histones can drive dramatic alterations in chromatin regulation and cellular identity, as exemplified by cancer-associated oncohistone mutations and germline mutations linked to neurodevelopmental and neurodegenerative disorders^4–7^. Similarly, mutagenesis studies in yeast have shown that single amino acid substitutions within the histone core can severely impact viability^3^. Histone variants further illustrate the sensitivity of chromatin function to subtle sequence variation. These non-allelic isoforms replace canonical histones within nucleosomes and can enable specialized chromatin states with minimal sequence differences^8–10^. A striking example is provided by the H2A.Z histone variant isoforms, H2A.Z.1 and H2A.Z.2, which differ by only three amino acids and present near-identical structures yet perform nonredundant biological functions^11–14^. Remarkably, their functional divergence can depend on a single residue within the buried histone core, serine or threonine at position 38^12^. Yet, given the lack of differences observed in nucleosome crystal structures, the molecular mechanisms by which subtle sequence variations are translated into large functional outcomes remain unclear.

Here we uncover a previously unrecognized layer of chromatin regulation mediated by the conformational plasticity of the nucleosome core. We show that subtle perturbations propagate allosterically across the nucleosome, altering its accessibility and recognition by chromatin-associated factors and ultimately influencing cell identity. These findings provide a unifying explanation for how subtle histone variations can be amplified into large biological effects, and further suggest that nucleosome core allostery represents a general principle of chromatin regulation.

### H2A.Z allosterically remodels nucleosome core dynamics

H2A.Z is a highly conserved variant of histone H2A that shares ∼60-65% sequence identity and a conserved histone fold^15,16^. Despite this overall conservation, H2A.Z presents an expanded acidic patch and sequence divergence within the C-terminal region, spanning both the H2A-H2B dimer docking domain and the C-terminal tail (**Fig.1A; Fig. S1A**). These differences are thought to tune nucleosome stability, turnover, and protein interactions^17^.

**Figure 1:**
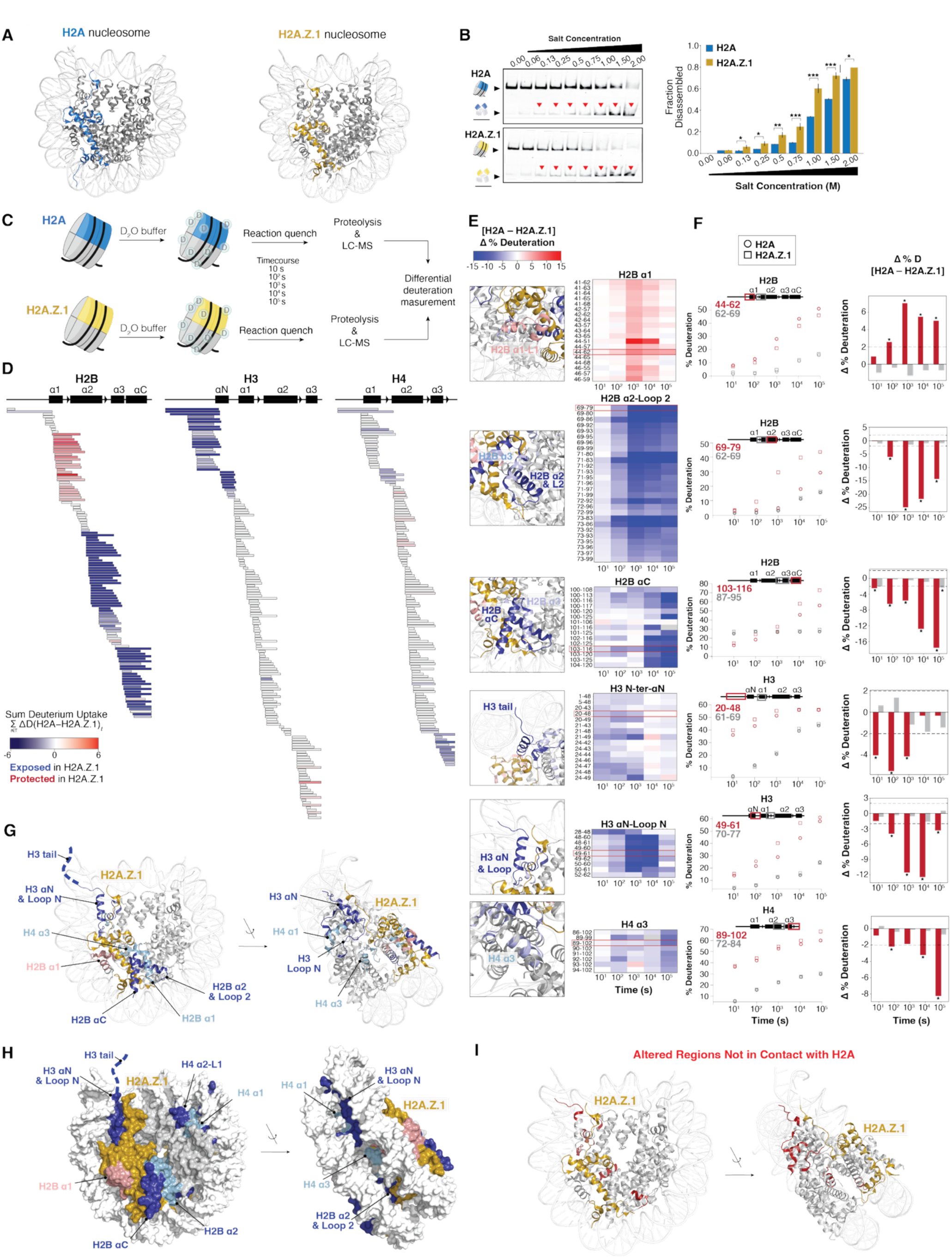
H2A.Z rearranges nucleosome core dynamics. A) Crystal structures of nucleosomes containing canonical H2A (blue, left panel, PDB:2CV5) or H2A.Z.1 (yellow, right panel, PDB: 5B33). For clarity, only one histone copy is colored; histones are shown in gray and DNA in white. B) Nucleosome salt-sensitivity assay. Left: representative FAM-fluorescence scan of a native gel shift assay of H2A (top) and H2A.Z.1 (bottom) nucleosomes exposed to increasing NaCl (0-2 M). Red triangles denote conditions with *P* < 0.005. Right: quantification of nucleosome dissociation, expressed as the percentage of total FAM-labeled DNA per lane. Error bars represent SEM (n=3); statistical differences (two-tailed t-test) are indicated. C) Schematic of the HDX-MS workflow. D) Sum of differential deuterium uptake (ΣΔD) between H2A- and H2A.Z.1-containing nucleosomes across all time points. Each bar represents a peptide, mapped below histone secondary structure elements and color-coded by percent difference in uptake. E) Left: mapping of differential regions onto the nucleosome structure. H2A.Z.1 is in yellow. Right: heat map of differential deuterium uptake (Δ%D) for selected histone regions across all time points. F) Left: deuterium uptake kinetics for representative differential peptides (red), with peptide locations shown in red boxes. For each peptide, a representative nondifferential peptide from histone regions showing comparable deuteration is shown for comparison (gray). Right: differential percent deuteration (Δ%D) plotted for each time point. The red dotted lines denote the significance threshold, calculated using a global pooled standard deviation with propagated error (see Tables S1-S2). G) Regions showing differential deuteration mapped onto the H2A.Z.1 nucleosome structure; colors indicate increased (blue) or decreased (red) accessibility. For clarity, only one histone copy is colored; DNA is white, H2A.Z.1 is yellow. H) Surface representation of panel G showing contacts between differentially deuterated regions and H2A. I) Differentially deuterated regions lacking direct H2A contacts (distance > 4 Å) highlighted in red.

To understand the mechanisms underlying the functional effects of H2A.Z incorporation, we examined the effects of this variant on nucleosome stability and conformational dynamics. We first evaluated nucleosome salt sensitivity by determining the ionic strength required for unwrapping DNA from human nucleosomes containing either canonical H2A or H2A.Z.1, the most abundant and widely studied H2A.Z isoform. Nucleosomes were exposed to increasing salt concentrations, and disassembly was quantified by monitoring the level of free DNA on native gels. H2A.Z.1 nucleosomes displayed greater salt sensitivity compared to H2A nucleosomes, with significantly higher disassembly across all salt concentrations tested (**Fig. 1B**), consistent with previous reports^13^, and indicating that variant incorporation alters nucleosome properties.

To dissect underlying conformational dynamics, we employed hydrogen–deuterium exchange mass spectrometry (HDX-MS), which reports on protein conformational dynamics via the exchange of backbone amide hydrogens with solvent deuterium at the level of individual peptides. Dual digestion with porcine pepsin and aspergillopepsin achieved >95% residue coverage, enabling analysis of all core histones (**Fig. S1B**; **Table S1**). We monitored deuterium incorporation in H2A and H2A.Z.1 nucleosomes over time (**Fig. 1C; Fig. S1C**). Differential uptake was used to quantify changes in backbone amide exchange, which reflect local conformational stability and structural accessibility within the nucleosome. Widespread differences were observed across all histones within the nucleosome, indicating substantial alterations in local structural dynamics or accessibility (**Fig. 1D-H; Fig. S1D, E**). These changes were localized to specific regions within each histone, with peptides spanning these sites showing consistent, significant differences in deuteration compared to peptides mapping to unaffected regions (**Fig. 1F, red vs gray**).

Histone H2B exhibited the most pronounced differences, with increased accessibility in H2A.Z nucleosomes at the C-terminal portion of the α2 helix and loop L2, the αC helix, and a modest decrease at the C-terminus of the α1 helix (**Fig. 1D-H; Fig. S1D, E**). The H2B αC and α1 helices are in contact with H2A/H2A.Z.1 and are surface-exposed regions known to interact with chromatin factors^18,19^. In contrast, the C-terminus of the H2B α2 helix and loop L2 are typically buried within the nucleosome core at the interface with the H3-H4 tetramer and DNA. These changes agree with recent reports of increased H2B dynamics within H2A.Z-H2B dimers^20^ and suggest loosening of both the dimer–DNA and dimer–tetramer contacts in H2A.Z.1 nucleosomes, aligning with the elevated salt sensitivity (**Fig. 1B**).

Remarkably, accessibility changes extended beyond the immediate H2A-H2B interface into distal and buried regions of H3 and H4 (**Fig. 1D-H; Fig. S1D-E**). In H3, the αN helix, loop LN, and adjacent N-terminal tail are more exposed in H2A.Z.1 nucleosomes. These elements, which anchor DNA at the dyad^1,21^, suggest weaker H3-DNA contacts and altered tetramer geometry, potentially influenced by proximity of the H2A.Z.1 C-terminus to the H3 αN helix^1,22^. Similarly, H4 showed increased deuterium uptake within the α3 helix-loop L3 region (**Fig. 1D-H**), and more modestly, the α1 helix at the later time points (**Fig. S1F-G**). These regions, which stabilize the H3–H4 tetramer and contact DNA, do not directly contact H2A/H2A.Z.1 (**Fig. 1H, I**), suggesting that perturbations introduced at the H2A.Z interface propagate allosterically throughout the nucleosome core to influence structurally distal and buried regions (**Fig. 1H, I**).

Overall, our HDX-MS reveals that histone variant incorporation does not simply alter accessibility at regions in direct contact with the variant, but instead allosterically reshapes the conformational landscape of the entire nucleosome, affecting both histone-DNA and histone-histone interfaces. These findings provide a mechanistic framework for how histone variants impair specific functions and suggest that distinct histone compositions may regulate genome function by controlling the accessibility of the entire nucleosome.

### H2A.Z isoforms allosterically encode distinct nucleosome core dynamics

Mammals express two H2A.Z isoforms, H2A.Z.1 and H2A.Z.2, encoded by the paralogous genes *H2AFZ* and *H2AFV*^10,11^. Both are ubiquitously expressed, with H2A.Z.1 generally predominant across tissues^24^. The isoforms differ by only three amino acids: residue 14 in the N-terminal tail, residue 38 in the globular core, and residue 127 in the C-terminal tail (**Fig. 2A; Fig. S2A**). Nucleosomes containing either isoform show nearly identical crystal structures, with a global Cα RMSD <0.3 Å; the only reported structural difference is a subtle shift in the L1 loop harboring residue 38^13^. Despite these minimal sequence and structural differences, the isoforms perform distinct, nonredundant roles in gene regulation, development, and disease, and each only partially rescues loss of the other^10,12,14,24–27^. How such minimal amino acid changes generate divergent biological outcomes remains unclear. Our HDX studies comparing H2A- and H2A.Z.1-containing nucleosomes raise the possibility that these subtle variant differences may induce distinct conformational dynamics that functionally regulate nucleosome accessibility.

**Figure 2:**
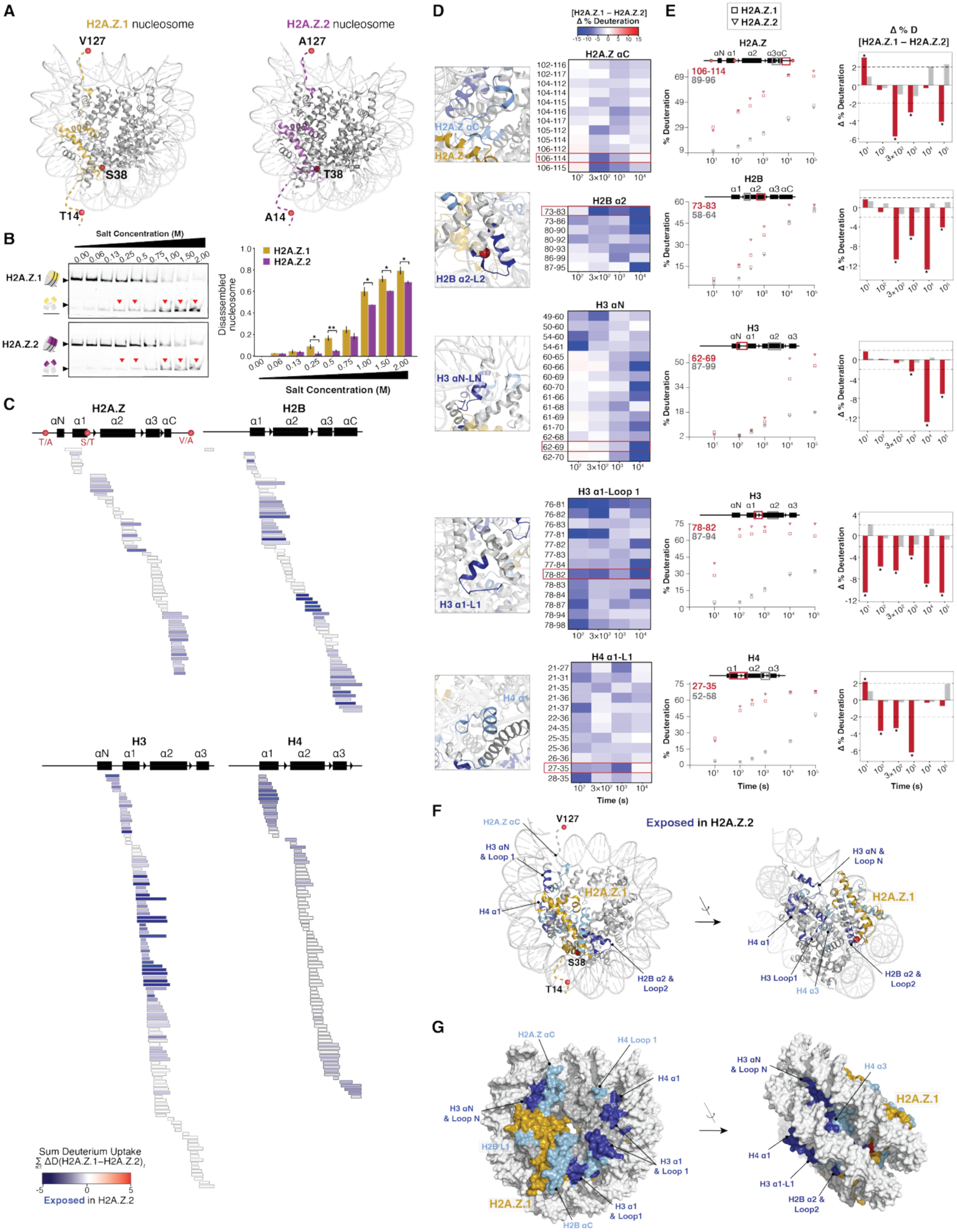
H2A.Z.1 and H2A.Z.2 nucleosomes exhibit distinct core conformational dynamics. A) Crystal structures of nucleosomes containing either H2A.Z.1 (yellow, left; PDB: 5B33) or H2A.Z.2 (purple, right; PDB: 3WAA). For clarity, only one copy of each histone is colored; histones are shown in light gray and DNA in white. B) Nucleosome salt-sensitivity assay. Left: representative FAM-fluorescence scan of a native gel shift assay of H2A.Z.1 (top) and H2A.Z.2 (bottom) nucleosomes exposed to increasing NaCl (0-2 M). Red triangles highlight conditions with *P* < 0.05. Right: quantification of nucleosome disassembly, expressed as the percentage of total FAM-labeled DNA per lane. Error bars represent the SEM (n = 3). Statistical differences (two-tailed t-test) are indicated. C) Sum of the differential deuterium uptake (ΣΔD) between H2A.Z.1- and H2A.Z.2-containing nucleosomes across all time points. Each bar represents a peptide, mapped below histone secondary structure elements and color-coded by percent difference in uptake. D) Left: mapping of differential regions onto the nucleosome structure. H2A.Z.1 is yellow. Right: heat map of differential deuterium uptake (Δ%D) for selected histone regions across all time points. E) Left: deuterium uptake kinetics for representative differential peptides (red), with peptide locations shown in red boxes. For each peptide, a representative nondifferential peptide from histone regions showing comparable deuteration is shown for comparison (gray). Right: differential percent deuteration (Δ%D) plotted for each time point. The red dotted lines denote the significance threshold, calculated using a global pooled standard deviation with propagated error (see Tables S1-S2). F) Histone regions exhibiting increased deuteration in H2A.Z.2 nucleosomes are mapped onto the H2A.Z.1 nucleosome structure in blue. The shade of blue reflects the magnitude of the deuteration increase. For clarity, only one histone copy is colored; DNA is white and H2A.Z.1 is yellow. G) Surface representation of regions with increased accessibility in H2A.Z.2 nucleosomes, as in panel F. Both histone copies are colored.

We first compared H2A.Z.1 and H2A.Z.2 nucleosomes by measuring salt-dependent disassembly. Nucleosomes containing either isoform were significantly more salt sensitive than canonical H2A nucleosomes (**Fig. S2B**). However, H2A.Z.2 nucleosomes were consistently more resistant to salt-induced disassembly than H2A.Z.1 nucleosomes across a range of ionic strengths (**Fig. 2B**), indicating that three amino acid substitutions are sufficient to alter nucleosome biophysical properties.

We then compared the HDX-MS profiles of the two nucleosomes. H2A.Z.2 nucleosomes exhibited increased solvent accessibility across all core histones relative to H2A.Z.1 (**Fig. 2C-G; Fig. S2C-F**). Thus, unlike the comparison between H2A.Z.1 and canonical H2A nucleosomes (**Fig. 1B**), the greater salt tolerance of H2A.Z.2 nucleosomes (**Fig. 2B**) correlated with increased nucleosome dynamics.

Because peptides spanning the substituted residues cannot be compared directly, HDX analysis was limited to flanking regions where coverage was available. Under these constraints, H2A.Z.2 nucleosomes showed a modest increase in accessibility within the H2A αC helix adjacent to V127A, whereas changes near S38T are smaller and less consistent (**Fig. 2C–E**). No peptide coverage was obtained for T14A (**Fig. 2C; Fig. S2D**). Remarkably, only a small subset of accessibility changes lay near the variant-specific residues (**Fig. 2E-G; Fig. S2D-F**). In H2B, the largest increase in accessibility was observed at the C-terminal end of the α2 helix and loop L2, which contact both the DNA backbone and H2A.Z loop L1, along with more modest changes in the αC helix and the buried α1-L1 region. In H3, enhanced solvent exposure mapped to the αN–loop LN regions positioned between the DNA gyre and the H2A.Z C-terminal tail, as well as the C-terminal end of the α1 helix and loop L1. H4 also showed increased deuteration in regions that are buried against the DNA, including the N-terminal end of the α1 and loop L1, and a modest increase in the deeply buried α3 helix. These HDX differences across multiple histone–DNA and histone–histone interfaces are consistent with localized conformational changes driven by point mutations, rather than global structural disruption. Notably, most perturbations occurred far from the substituted residues and in structurally connected regions, suggesting that the three amino acid differences propagate long-range allosteric rearrangements that tune the nucleosome conformational landscape rather than inducing a new structure or global unfolding.

Our results demonstrate that three residue differences within H2A.Z are sufficient to remodel nucleosome conformational dynamics through long-range allosteric effects, providing a mechanistic framework for how nearly identical H2A.Z isoforms carry out distinct chromatin functions. More broadly, these findings establish a generalizable mechanism by which minimal sequence variation within histones can encode distinct nucleosome conformational dynamics that allosterically modulate nucleosome accessibility beyond what static structures reveal.

### A single buried substitution allosterically reprograms nucleosome core dynamics

Among the three amino acid differences between H2A.Z.1 and H2A.Z.2, T14A and V127A lie in the flexible N- and C-terminal tails, whereas S38T resides within the globular core (**Fig. 2A**). To isolate the contribution of the core residue, we generated a chimeric histone consisting of H2A.Z.1 tails (T14 and V127) and the H2A.Z.2 core (T38) (**Fig. 3A**). At low salt concentrations (<0.5 M NaCl), nucleosomes containing the chimera behaved similarly to H2A.Z.2, whereas at higher salt (1–2 M NaCl) they showed an intermediate phenotype, indicating that threonine 38 contributes to the increased salt stability of H2A.Z.2 nucleosomes but is not sufficient to fully recapitulate the H2A.Z.2 phenotype (**Fig. 3B**).

**Figure 3:**
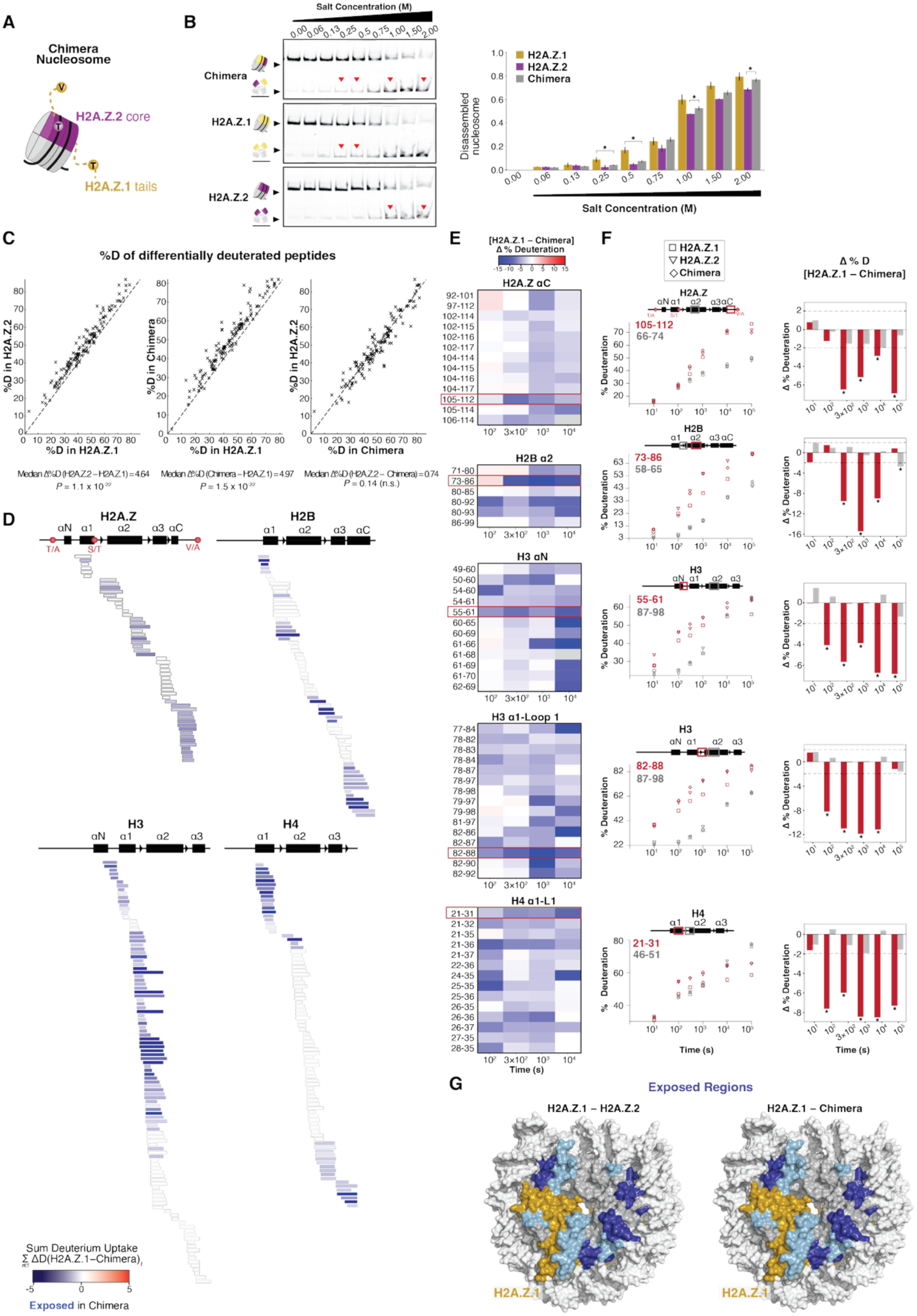
Core residue 38 regulates nucleosome core dynamics. A) Cartoon illustration of the chimeric nucleosome containing H2A.Z.1 tails and the H2A.Z.2 core. B) Nucleosome salt-sensitivity assay. Left: representative FAM-fluorescence native gel-shift assays of the indicated nucleosomes exposed to increasing NaCl concentrations (0-2 M). Red triangles highlight conditions with *P* < 0.05. Right: Quantification of nucleosome disassembly, expressed as the percentage of total FAM-labeled DNA per lane. Error bars represent the SEM (n = 3); statistical differences (two-tailed t-test) are indicated. C) Correlation plots of deuterium uptake (%D) for all differentially deuterated peptides between H2A.Z.1 and H2A.Z.2, H2A.Z.1 and Chimera, or H2A.Z.2 and Chimera. Each cross represents one peptide’s %D in one nucleosome (x-axis) versus the other (y-axis). The diagonal line denotes equal deuteration in the compared nucleosomes. The median difference and one-sided Wilcoxon test for each comparison are indicated, showing that these differential peptides are significantly more deuterated in H2A.Z.2 and the Chimera than H2A.Z.1. Nondifferential peptides (not shown) showed no significant difference across all nucleosome pairs (P > 0.05). D) Sum of the differential deuterium uptake (ΣΔD) between H2A.Z.1- and Chimera-containing nucleosomes across all time points. Each bar represents a peptide, mapped below histone secondary structure elements and color-coded by percent difference in uptake. E) Heat map of differential deuterium uptake (Δ%D) for selected histone regions across all time points. F) Left: deuterium uptake kinetics for representative differential peptides (red), with peptide locations shown in red boxes. For each peptide, a representative nondifferential peptide from histone regions showing comparable deuteration is shown for comparison (gray). Right: Differential percent deuteration (Δ%D) plotted for each time point. The red dotted lines denote the significance threshold, calculated using a global pooled standard deviation with propagated error (see Tables S1-S2). G) Histone regions exhibiting increased deuteration in H2A.Z.2 and in the Chimera relative to H2A.Z.1 are mapped in blue on the surface representation of the nucleosome. The shade of blue reflects the magnitude of deuteration increase. Both histone copies are colored. DNA is white and H2A.Z.1 is yellow.

To determine whether residue 38 similarly modulates nucleosome dynamics, we performed HDX-MS on the chimera (**Fig. 3C-F; Fig. S3A**). If the core residue 38 is a major contributor to isoform-specific changes, the chimera should exhibit a deuteration pattern more similar to H2A.Z.2 than to H2A.Z.1. Across all differentially deuterated peptides, the chimera indeed tracked with H2A.Z.2 and diverged from H2A.Z.1 (**Fig. 3C**). Analysis of the absolute per-peptide differences in deuterium uptake further showed that both H2A.Z.2 and the chimera displayed significantly larger median differences from H2A.Z.1 than from each other (**Fig. S3B**), confirming that the chimera recapitulates the H2A.Z.2 conformational landscape and that residue 38 is the dominant contributor to the isoform-specific conformational changes.

Consistent with this, regions of increased accessibility in the chimera overlapped with those altered in H2A.Z.2 nucleosomes (**Fig. 3D-F; Fig. S3C-E**), including the H2A.Z αC helix; the H2B α2 helix-loop L2; the H3 αN helix–loop LN and α1 helix-loop L1; and the H4 α1 helix-loop L1 and the N-terminal portion of the α2 helix. These shared changes indicate that the S38T substitution alone is sufficient to recapitulate most of the long-range rearrangements observed in H2A.Z.2 nucleosomes.

Together, these findings reveal that a single conservative amino acid substitution within the H2A.Z core, S38T, is sufficient to modulate salt sensitivity and allosterically reprogram the conformational landscape of the nucleosome. Located within the α1 helix at the interface of both histone–DNA and histone–histone contacts, residue 38 acts as a central regulatory residue that exerts long-range allosteric effects propagating to distal nucleosome regions. More broadly, these results establish that even minimal perturbations within the nucleosome core can allosterically encode distinct conformational dynamics, such that the effects of a single residue change are amplified across the entire nucleosome to alter its surface accessibility.

### Nucleosome dynamics control chromatin factor recruitment

Conformational changes within the nucleosome can expose normally buried surfaces or alter surface accessibility, thereby modulating both inter-nucleosome interactions and recognition by chromatin factors. To assess consequences for higher-order chromatin contacts, we performed *in vitro* phase separation assays in which chromatin fibers demix into condensed droplets through inter-nucleosomal interactions mediated by both histone cores and tails^28,29^. We assembled 12-nucleosome arrays containing canonical H2A, H2A.Z.1, H2A.Z.2, or the H2A.Z chimera. Chromatin fibers containing either H2A.Z isoform required higher threshold concentrations to phase-separate than those containing canonical H2A (**Fig. 4A; Fig. S4A**), indicating that incorporation of H2A.Z reduces the propensity of nucleosome arrays to engage in the multivalent interactions necessary for condensate formation. Notably, H2A.Z.1 arrays displayed a two-fold higher phase separation threshold concentration than H2A.Z.2 arrays, indicating that three amino acid substitutions substantially alter the inter-nucleosome interactions driving condensation (**Fig. 4A**). To dissect tail vs core contributions, we examined chimera arrays containing H2A.Z.1 tails and the H2A.Z.2 core. The phase separation threshold of the chimera matched that of H2A.Z.1 (**Fig. 4A**), indicating that the tail residues, rather than the core, primarily govern inter-nucleosomal interactions.

**Figure 4:**
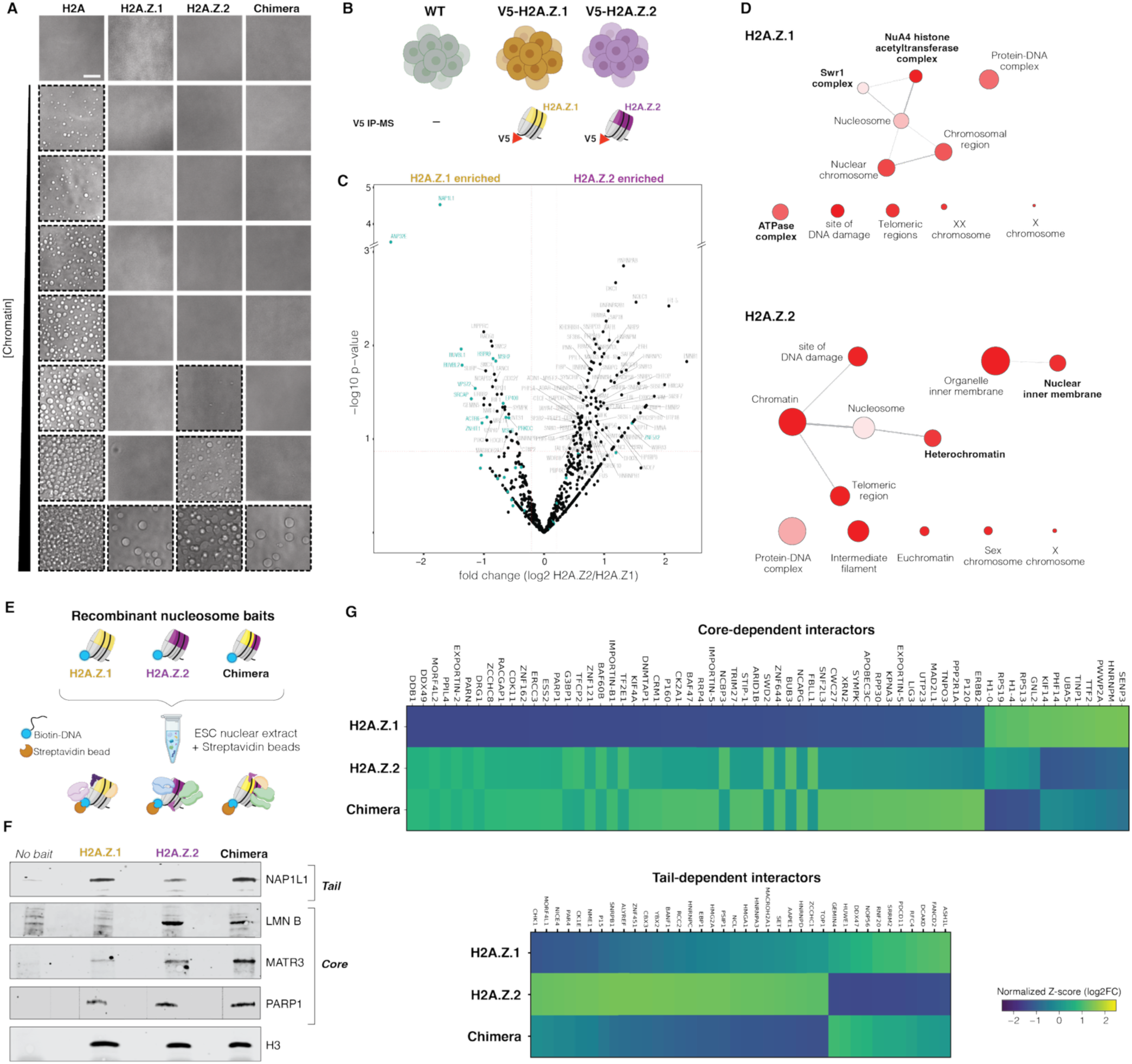
H2A.Z core and tails differentially regulate chromatin compaction and interaction networks. A) Phase separation assays of 12-nucleosome arrays containing H2A, H2A.Z.1, H2A.Z.2, or chimera nucleosomes across increasing chromatin concentrations under physiological salt conditions. Dotted boxes indicate conditions in which condensates are observed. Scale bar, 4 μm. B) Schematic of the V5-tagged IP-MS strategy in hESCs endogenously expressing V5-tagged H2A.Z.1 or H2A.Z.2. C) Volcano plot showing proteins differentially enriched in H2A.Z.1 versus H2A.Z.2 V5 IP-MS conditions. Known interactors are highlighted in teal. D) Cytoscape Gene Ontology (GO) Cellular Component network of proteins enriched in H2A.Z.1-(top) or H2A.Z.2-(bottom) V5 IP-MS. Node size reflects the number of proteins shared between GO terms. E) Schematic of the biotinylated nucleosome pull-down assay. F) WB of streptavidin pull-downs showing enrichment of the indicated interactors. G) Heat map of factors identified by MS, clustered according to core- or tail-dependent recognition.

We next asked whether nucleosome dynamics influence recognition by chromatin factors. To identify isoform-specific interactors in cells, we performed V5 immunoprecipitation followed by TMT-based quantitative mass spectrometry (IP-MS) in human embryonic stem cells (hESC) expressing endogenously tagged H2A.Z isoforms (**Fig. 4B**)^12^. Because H2A.Z.1 is approximately five-fold more abundant than H2A.Z.2, we used an excess of V5-H2A.Z.2 extract to normalize H2A.Z input and achieve comparable pull-down efficiency (**Fig. S4B**). We recovered known H2A.Z binders, including the H2A.Z chaperones NAP1L1 and ANP32E, and the remodelers SRCAP and the INO80 component RUVBL1/2^15,30–36^ (**Fig. 4C; Fig. S4C-D**). Several candidates were further validated by IP western blotting (WB) (**Fig. S4E**). H2A.Z.1 co-purified with chromatin remodelers (SRCAP, RUVBL1/2, SMARCA5/6, HELLS) and DNA-repair factors (RAD50), whereas H2A.Z.2 associated with nuclear-envelope proteins (LAMIN A/B, LBR, MATR3) and silencing factors (CBX1, CBX3, CBX5, DNMT3) (**Fig. 4D**), suggesting that distinct nucleosome conformational states preferentially recruit different chromatin regulators.

To assess isoform-specific recognition without confounding cellular factors and dissect tails vs core contributions, we performed *in vitro* pull-downs using biotinylated mono-nucleosomes as bait (**Fig. 4E; Fig. S4F**) and probed for the identified H2A.Z interactors. NAP1L1 preferentially bound H2A.Z.1 nucleosomes and retained comparable affinity for the chimera, indicating tail-dependent recognition (**Fig. 4F; Fig. S4G**). In contrast, linker histone H1 showed tighter binding by fluorescent polarization to H2A.Z.1 than H2A.Z.2 and the chimera, indicating core-dependent modulation of H1 association (**Fig. S4H).** LAMIN B and MATR3, and to a lesser extent PARP1, were enriched on both H2A.Z.2 and chimera nucleosomes, revealing that the S38T core substitution enhances the binding of these factors (**Fig. 4F; Fig. S4G**).

To systematically assess how tail and core substitutions shape H2A.Z interactomes, we performed TMT-based quantitative MS on pull downs using all three nucleosome baits. Comparison of chimera interactors with those of each isoform revealed two classes of interactors: core-dependent factors, whose binding mirrored H2A.Z.2 (**Fig. 4G; Fig. S4I)**, and tail-dependent factors (**Fig. 4G; Fig. S4J)**, whose binding tracked with H2A.Z.1 nucleosomes. Notably, because residue 38 is buried and not solvent-accessible, core-dependent interactions must arise from allosterically driven changes in nucleosome conformation that alter its surface accessibility.

Together, these results reveal that the three amino acids distinguishing H2A.Z.1 and H2A.Z.2 modulate both inter-nucleosome interactions and chromatin factor recruitment. Whereas chromatin folding is primarily regulated by H2A.Z tail identity, the interactome reflects contributions from both tails and core residues. More importantly, the interactions regulated by the substitution at residue 38 show that a single buried residue can control factor recognition by allosterically reshaping the nucleosome surface. This suggests that nucleosome conformational dynamics amplify local perturbations within the nucleosome core into global changes that regulate factor recruitment. More broadly, these findings establish that nucleosome core dynamics encode regulatory information that is read out by chromatin factors, providing a unifying mechanism by which subtle perturbations within the nucleosome core are amplified to control genome function.

### Allosteric core residue 38 regulates stem cell state transitions

H2A.Z is essential for development and differentiation^15^. In embryonic stem cells, H2A.Z is enriched at promoters and enhancers of lineage-specific genes, where it contributes to a poised chromatin state that enables rapid transcriptional activation upon differentiation^37,38^. Loss of H2A.Z disrupts this balance, impairing chromatin accessibility and lineage commitment^8,10,38,39^. H2A.Z has been implicated in both gene activation and repression, yielding seemingly contradictory models for its function^16,40^. These inconsistencies suggest that H2A.Z’s effects are highly context-dependent, influenced by genomic location, interacting partners, and potentially isoform-specific conformational dynamics.

To determine the functional consequences of altering nucleosome conformational dynamics, we used CRISPR/Cas9 to edit the endogenous *H2AFZ* and *H2AFV* loci in H9 hESCs, swapping Ser38 and Thr38 of each isoform (**Fig. 5A; Fig. S5A**). This strategy preserves endogenous isoform abundance, avoiding dosage effects, and leaves all surface-exposed histone residues unchanged, thereby isolating effects arising specifically from changes in nucleosome conformations rather than altered nucleosome surfaces. Because our HDX-MS revealed that a single-residue substitution can profoundly alter nucleosome properties, we avoided epitope tagging to ensure an uncompromised assessment of residue 38 function.

**Figure 5:**
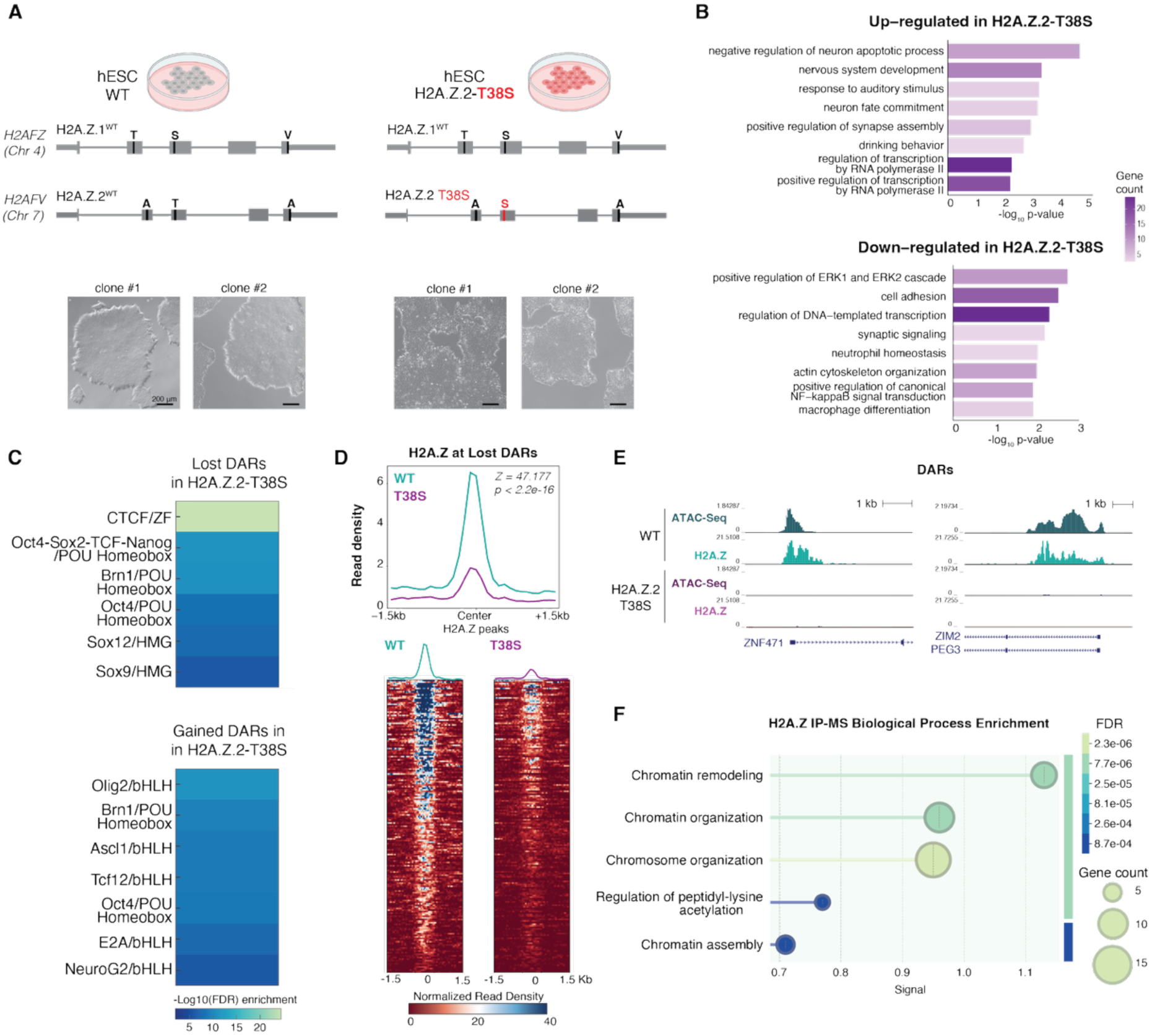
H2A.Z.2 T38S mutation biases cell state. A) Top: Schematic of the genotypes of WT and H2A.Z.2-T38S hESC lines generated in this study. Bottom: Bright-field images of two WT and two mutant clones. Scale bar, 200 μm. B) Gene Ontology categories enriched among genes upregulated (top) and downregulated (bottom) in H2A.Z.2-T38S cells relative to WT. C) Motif enrichment analysis for DARs that lose (top) or gain (bottom) accessibility in H2A.Z.2–T38S cells. Motifs are ranked by enrichment significance (–log10 FDR). D) Top: Metaplot of H2A.Z CUT&Tag signal centered on ATAC-seq peaks that lose accessibility in H2A.Z.2-T38S cells. Z-score and permutation test *P* value are indicated. Bottom: Heat map showing H2A.Z CUT&Tag signal at regions with reduced accessibility in H2A.Z.2-T38S cells. E) Representative genome browser snapshots illustrating loss of H2A.Z signal at lost DARs. F) Functional enrichment analysis of H2A.Z-interacting proteins differentially enriched between WT and H2A.Z.2-T38S cells, as identified by IP-MS. Terms are ranked by FDR.

Despite multiple attempts, we were unable to recover H2A.Z.1-S38T clones, suggesting that Ser38 is essential for H2A.Z.1 function, consistent with the lethality of *H2AFZ* loss in mouse embryos and mouse ESCs^37,41,42^. In contrast, H2A.Z.2-T38S clones were readily obtained (**Fig. S5A**), and two independent clones were used for downstream analysis.

H2A.Z.2-T38S cells exhibited morphological changes. Unlike wild type (WT) hESCs, which form smooth, compact colonies, mutant colonies displayed jagged borders and a dispersed appearance (**Fig. 5A**), indicating changes in cell state and spontaneous differentiation.

Transcriptomic profiling revealed that H2A.Z.2-T38S cells spontaneously initiate neural lineage commitment (**Fig. 5B; Fig. S5B-C**). Pluripotency regulators (e.g. ESRG, DPPA2, DPPA3, SOX3, PEG3), cell-cycle genes (STAG2, CCNB3, NSMCE1), and adhesion factors (ICAM3, CD44, VCAN, TAGLN, multiple PCDH genes) were downregulated (**Fig. 5B**). Conversely, neurogenic transcriptional programs and lineage-specifying factors were activated (**Fig. 5B; Fig. S5C**), consistent with the established role of H2A.Z.2 in neural pathways^14,24,34,43^.

ATAC-seq profiling revealed changes in chromatin accessibility (**Fig. S5D-F**). Regions losing accessibility were enriched for pluripotency-associated motifs such as OCT4, SOX2, and NANOG, as well as the architectural protein CTCF (**Fig. 5C**), suggesting a coordinated silencing of stem cell regulatory elements. Newly accessible regions in H2A.Z.2-T38S cells were enriched for basic helix–loop–helix (bHLH) transcription factor motifs, including core regulators of neural lineage specification such as Olig2, Ascl1, NeuroG2, and Tcf12 (**Fig. 5C**). These accessibility changes mirror the transcriptional reprogramming observed in RNA-seq (**Fig. 5B**) and indicate a shift toward a neural-primed state.

Both H2A.Z.1 and H2A.Z.2 isoforms are enriched at promoters and regulatory elements and generally co-occur genome wide (>70% of shared peaks) (**Fig. S5G-I**). Genome-wide profiling of total H2A.Z revealed that occupancy was largely preserved in H2A.Z.2-T38S mutant, with >80% of peaks shared between WT and mutant cells (**Fig. S5J**). While no global changes in H2A.Z isoforms gene expression were observed (**Fig. S5K)**, a modest reduction in H2A.Z chromatin signal was detected in the mutant (**Fig. S5L**), consistent with reported decreases in H2A.Z levels upon differentiation^44,45^.

Redistribution of H2A.Z was most pronounced and significantly enriched at differentially accessible regions (DARs) (*p* = 3.3 e-78, Z-score = 18.684). Sites losing chromatin accessibility exhibited significantly reduced H2A.Z signal (*p* < 2.2 e-16, Z-score = 47.177) (**Fig. 5D**). This is consistent with previous studies linking H2A.Z to increased chromatin accessibility and indicates that altered H2A.Z localization contributes to increased chromatin accessibility in H2A.Z.2-T38S mutants^10,38,46,47^. Conversely, sites gaining chromatin accessibility did not show consistent changes in H2A.Z occupancy (*p* = 0.99, Z-score = 3.593), suggesting that they do not simply arise from increased H2A.Z incorporation (**Fig. S5M, N**).

We then asked whether mutation of residue 38 alters the H2A.Z interactome. H2A.Z IP-MS revealed a loss of multiple factors associated with H2A.Z-related pathways in the H2A.Z.2-T38S cells, including lamina-associated proteins (LMNB2, BAF, H1), and DNA repair factors (XRCC5, XRCC6, FANCI, FANCD2) (**Fig. 5F, Fig. S5O**). Several chromatin remodelers were also differentially associated, including SMARCB1 and SMARCC1, core subunits of the SWI/SNF complex; SNF2H (SMARCA5) and HELLS; and CHD4, the ATPase subunit of the NuRD complex. Additional chromatin remodeling and nucleosome regulatory factors were also affected, including the NuRD components MTA3 and RBBP4, as well as the histone chaperones NAP1L1 and ANP32E. Thus, the T38S core substitution reshapes H2A.Z interactions, likely reflecting both altered direct nucleosome recognition and indirect effects arising from altered chromatin context.

Together, these results show that the buried T38S substitution alters chromatin accessibility and factor recruitment, resulting in genome-wide changes in gene expression and altered cell identity. The buried location of residue 38 provides a direct link between allosteric remodeling of nucleosome accessibility and functional chromatin changes. More broadly, these findings establish the functional importance of nucleosome core dynamics as an additional layer of chromatin regulation, through which subtle perturbations are amplified into global cellular outcomes.

## Discussion

Our findings support a model in which the nucleosome core functions as an allosteric regulatory module that governs chromatin function through its conformational landscape. By integrating HDX-MS with functional and biochemical analyses, we show that subtle perturbations within the histone core propagate across the nucleosome to control accessibility, chromatin factor recognition, and ultimately cell identity. These results indicate that nucleosome allostery can amplify minimal molecular variation into substantial regulatory outcomes, even in the absence of detectable changes in static structure, thereby establishing nucleosome core dynamics as a fundamental axis of chromatin regulation.

### Allosteric regulation within the nucleosome core

Chromatin regulation has traditionally been interpreted through the lens of histone tail modifications, which serve as signaling platforms, while the globular core has been largely viewed as a structural scaffold^2^. Our results challenge this tail-centric paradigm by showing that the nucleosome core itself encodes regulatory information embedded within its intrinsic conformational landscape. Small perturbations within the core propagate allosterically across histone–DNA and histone–histone interfaces, reshaping nucleosome accessibility and interaction surfaces far from the site of perturbation. In this context, local changes are transmitted through structurally coupled regions to regulate global accessibility. This mode of regulation resembles that of well-established allosteric systems, such as GPCRs, in which function is controlled by shifts in conformational ensembles rather than static structures^48^.

A striking consequence of these allosteric rearrangements is that even a single buried amino acid substitution, such as residue 38 in H2A.Z, is sufficient to globally reorganize histone–histone and histone–DNA contacts. By altering both the accessibility of specific interaction sites and the broader reorganization of interaction surfaces, these changes regulate chromatin factor binding. This provides a mechanistic explanation for how nucleosomes with nearly identical static structures can nonetheless encode distinct biological effects.

While previous studies have shown that chromatin regulators can induce conformational changes upon binding^28,49–52^, our findings demonstrate that nucleosome composition alone is sufficient to encode distinct conformational states. This shifts nucleosome plasticity from a responsive property to an intrinsic mechanism of regulation. The partial overlap between regions differentially accessible in H2A.Z variants and those remodeled upon factor binding^28^ further suggests the existence of allosteric hotspots that act as control nodes for chromatin regulation.

We propose that the nucleosome core acts as an allosteric integrator that translates local molecular variation into global regulatory outputs. In addition to histone variations, other sources of nucleosome perturbation, including post-translational modifications within the core and DNA sequence variation, may similarly reshape the nucleosome conformational landscape to regulate genome function. Thus, nucleosome allostery may provide a unifying principle for how multiple nucleosome features are functionally integrated.

### Nucleosome conformational landscape in chromatin recognition and cell identity

Our findings establish a direct link between nucleosome core dynamics and cell state transitions. We show that conformational changes within the nucleosome are functionally amplified through chromatin factor recruitment.

These results point to a fundamentally expanded model of chromatin recognition in which specificity is also determined by conformational dynamics that regulate the exposure of nucleosome binding interfaces. Chromatin factors can sense local changes in epitope accessibility that arise from long-range allosteric effects. In this way, core-encoded conformational states are translated into distinct interaction landscapes, enabling chromatin factors to discriminate between nearly identical static nucleosome structures. Remarkably, this implies that even buried residues can impact nucleosome recognition by allosterically reshaping epitope accessibility, nucleosome structure, and interaction surfaces without being directly contacted by binding factors. Such conformational selectivity provides a mechanism for integrating multiple inputs, including histone variants, core-localized perturbations, tail modifications, and DNA sequence, to specify factor recruitment.

### Broader functional implications

In disease, cancer-associated oncohistone mutations^4,5^ and mutations linked to neurodevelopmental and neurodegenerative disorders^6,7^ are frequently located within the histone core. Similarly, in development and evolution, histone sequences can differ by only a few core amino acids and yet drive functional diversification without detectable changes in static structures^8,10^. Despite these observations across diverse biological contexts, a unifying mechanistic understanding of how such minimal perturbations give rise to distinct regulatory outcomes has remained unclear, and nucleosome allostery may provide a framework to explain the underlying mechanisms.

Understanding how nucleosome dynamics are encoded and interpreted therefore has broad implications for development and disease, and may enable new therapeutic strategies aimed at targeting nucleosome conformational states. Mapping allosteric hotspots, defining how these states intersect with post-translational modifications, and identifying which chromatin factors preferentially recognize specific nucleosome conformations will be critical for elucidating how chromatin information is encoded and can be manipulated.

## Acknowledgements

We thank members of the Sanulli laboratory for helpful discussions and suggestions; N. Neff and the rest of the Stanford Chan Zuckerberg Biohub Center team for ongoing support; the Wysocka lab for providing V5-H2A.Z.1 and V5-H2A.Z.2 cell lines; and Hayden S Saunders for providing recombinant H1. AI-assisted tools were used for language editing and clarity; all scientific content and conclusions are the responsibility of the authors. This work utilized bioinformatics services and computing resources provided by the Stanford Genetics Bioinformatics Service Center (GBSC). We thank G. Narlikar, JD Gross, and R Bonasio for feedback on the manuscript.

## Author contributions

SS, ANW and MP conceived the project. ANW designed, performed, and interpreted the performed salt sensitivity assays. HDX-MS experiments were designed by ANW, DNK, SM, and SS, and performed by ANW and DNK. MP designed, performed, and interpreted IPs. MT performed and analyzed MS. MMW performed RNA-seq and Cut&Tag. ANW and MMW cultured H9 cells. Cell editing was performed by EditCo and bioinformatic analyses by the Genetics Bioinformatic Service Center and MMW. AN and JM prepared chromatin, performed FP, and phase separation assays. SS supervised the project and wrote the bulk of the manuscript with input from the authors. All authors discussed the results and commented on the manuscript.

## Materials and Methods

### Protein expression, purification and isotope labeling

All the histones were expressed and purified from *E. coli* following published protocols ^53^. All the mutant versions of the histones were made by quick-change site-directed mutagenesis. All mutations were confirmed by nanopore whole-plasmid sequencing.

### Mononucleosome Assembly

Histone octamer was assembled from purified histones by salt dialysis as described ^28^. The 601 positioning sequence (147 bp) DNA fragment was made using restriction enzyme digestion of a plasmid carrying multiple copies of 601 DNA fragment as described previously ^53^. 5’-FAM and 5’-biotin+20bp linker DNA were generated by PCR amplification using a 5’-labeled primer (IDT) followed by sizing column purification. Nucleosomes were assembled using published gradient dialysis-based protocols^53^. The purification of the nucleosomes was carried out on a 10 to 30% glycerol gradient or by gel filtration purification with a Superdex-200 column.

### Salt Sensitivity Assay

Nucleosomes assembled with 5’ FAM-labeled Widom 601 core DNA (147bp) were incubated at 25C for 1 hr at 200nM in 50mM HEPES (pH 7.5) buffer with 100 ng/ul stop plasmid and 0.065, 0.13, 0.25, 0.5, 0.75, 1, 1.5, or 2 M NaCl. After incubation, one fourth volume of loading buffer (20% glycerol, 50 mM HEPES pH 7.5) was added to each sample before running samples with non-denaturing PAGE (6% acrylamide, 0.5x TBE) at 120V for 2hrs. Gels were then imaged with Typhoon variable mode imager (GE Life Sciences, Pittsburgh, PA) by scanning for fluorescent labels. Assembled nucleosome and free DNA bands were then quantified by densitometry using ImageJ. The fraction of free DNA was determined by the ratio of free DNA to nucleosome normalized to the ratio from the no salt condition. At least three biological replicates were analyzed.

### HDX-MS

Each HDX experiment was carried out with biological replicate (see Table S1-2). Hydrogen exchange, protease digestion and LC-MS were performed as previously described^54^. Briefly, for each timepoint, 2 μl of >8uM nucleosomes were mixed with 8 μL of D_2_O-containing HDX buffer (20mM HEPES, pH 7.5; 150mM KCl; 10mM DTT, pD 7.6) and incubated at 25°C for a range of time points (0s, 10s, 10^2^s, 3×10^2^s, 10^3^s, 10^4^s or 10^5^s). Deuteration was quenched by adding 4°C quench buffer (0.5M TCEP, pH 2.5; 8M urea; 10% glycerol) at 1:1 ratio to the exchange reaction, followed immediately by flash freezing samples in liquid nitrogen. All samples were thawed at RT immediately before injection into a cooled valve system (Tajan LEAP) coupled to a LC (Thermo UltiMate 3000).The quenched sample was digested in-line at 200ul/min by two immobilized acid protease columns, first aspergillopepsin (Sigma-Aldrich P2143) and then porcine pepsin (Sigma-Aldrich P6887), maintained at 10°C prepared as described previously^54^. The resulting peptides were desalted on a hand-packed trap column (thermos Scientific POROS R2 reversed-phase resin 1112906, 1mm ID x 2cm, IDEX C130B) and then loaded onto a Waters Aquity analytical column (Cat. #186002344), both maintained at 2°C. The bound peptides were gradient-eluted (5–40% CH_3_CN w/v and 0.1% w/v formic acid) across the analytical column at 45ul/min for 16 min at 2°C and analyzed directly using a high-resolution Orbitrap mass spectrometer (Q Exactive, Thermo Fisher) in positive mode with settings for MS and MS/MS runs described previously^54^. After each run, the sample loop, protease and trap columns were manually washed at 200ul/min first with 200ul of guanidinium wash buffer (1.6 M GdmCl, 0.1% formic acid, pH 2.4), then with 200 ul of urea wash buffer (3.2M urea 0.6% formic acid, pH 2.7). Peptide identification was performed using Byonic (Protein Metrics) with the sequence of all histones forming the search library. For each exchange experiment, MS/MS were performed for each nucleosome type, and these peptide libraries were combined for downstream analysis. Peptic digestion of nucleosomes yielded >95% sequence coverage of the histone with 100% coverage across folded nucleosome regions and reduced coverage on the histone tails. Peptide isotope distributions at each time point were fit in HDExaminer 3 through automatic unimodal and bimodal peak calling followed by manual verification for consistency with the experimental spectra. Deuteration levels were determined by subtracting mass centroids of deuterated peptides from undeuterated peptides. Some peptides displayed bimodal spectra, where the less deuterated distribution was consistent with carryover between runs; accordingly, the more deuterated distribution was used to quantify deuterium uptake for these peptides. Percent deuterium uptake for each peptide was calculated using the theoretical maximum uptake for that peptide assuming an 80% maximum exchange, matching the deuterium concentration in the sample after 1:5 dilution in D_2_O HDX buffer. No correction was performed for back-exchange. Peptides containing the substituted residues were excluded from differential HDX analysis to avoid sequence-dependent effects on exchange and peptide behavior. Deuterium uptake for each peptide is calculated for each time point and the difference in %D and #D values between the nucleosomes is shown as a heat map with a color code given at the bottom of the figure. Thresholds of significance for difference in deuteration were determined for each data set as reported in Table S1-2.

### 12 Nucleosome-array Assembly

DNA was generated by restriction enzyme digestion of a plasmid containing 12 consecutive 601s spaced by 25 bp, followed by gel purification. The reconstitution of nucleosome arrays followed the protocols previously described^28^. Histone octamers are combined in equimolar amount with 12-mer DNA (12 repeats of the 601 DNA sequence separated by 25-bp linkers). Final dialysis against 20 mM HEPES pH 7.6, 1mM DTT was performed overnight. After assembly, arrays were usually used for experiments within 1–2 days after assembly. Since the 601 repeats in the 12-mer DNA sequence are separated by BstXI restriction enzyme sites, quality of assembly was assessed by BstXI digestion followed by native gel. Over-assembly was avoided by ensuring that >95% of the digested fragments migrated as mononucleosomes rather than slower migrating species. We note here that the quality of the arrays used in these studies had to be carefully controlled to avoid over assembly of histone octamers, as we noticed that over-assembled arrays displayed aberrant and non-reproducible aggregation behavior (**Fig. S4A**).

### Fluorescence Polarization

Nucleosome polarization assays were conducted in buffer containing 0.02% NP-40, 175 mM KCl, 20 mM HEPES pH 7.5, 1 mM DTT at 22°C. Each anisotropy sample contained a final nucleosome concentration of 2.5 nM and the H1.4 concentration was varied. The reaction was incubated 30 min at 22°C and fluorescence polarization was measured on a PHERAstar FSX (BMG Labtech). Data points from three independent H1.4 dilution curves were averaged and standard errors calculated. The following binding model was used to derive K_d_ Y=((X^ N*FPmax)+(Kd^ N*FPmin))/(X^ N+Kd^ N) in which Y is the fluorescence polarization signal observed; X is H1 concentration; FPmin is the fluorescence polarization signal for the probe alone; FPmax is the fluorescence polarization signal at saturating protein concentration; N is the Hill coefficient. Human histone H1.4 was kindly provided by the Narlikar lab.

### Cell Culture

Human H9 embryonic stem cells (WiCell WA09) were grown in mTeSR plus media (STEMCELL Technologies 100-0276) with Cultrex coating, following manufacture instructions.

### H9 *H2AFV* T38S Editing

CRISPR Cas9 mediated knock in cells of *H2AFV* T38S in H9 cells were generated by EditCo Bio, Inc. (Redwood City, CA, USA). Ribonucleoproteins containing the Cas9 protein and synthetic chemically modified guide RNA were electroporated into the cells along with a single-stranded oligodeoxynucleotide (ssODN) donor using EditCo’s optimized protocol. Editing efficiency is assessed upon recovery, 48 hours post electroporation. Genomic DNA is extracted from a portion of the cells, PCR amplified and sequenced using Sanger sequencing or NGS. The resulting Sanger chromatograms are processed using EditCo’s Inference of CRISPR edits software (https://ice.editco.bio/#/) or NGS bioinformatics pipeline. To create monoclonal cell populations, edited cell pools are seeded at 1 cell/well using a single cell printer into 96 or 384 well plates. All wells are imaged every 3 days to ensure expansion from a single-cell clone. Clonal populations are screened and identified using the genotyping strategy described above. Each clone was further validated using PCR-nanopore sequencing.

### Nucleosome Biotin IP-MS

Each IP was performed in triplicate using ∼1.6×10^7^ cells for each IP. Single cell suspension was obtained by using Accutase treatment (Millipore SCR005) and cells were rinsed in PBS 1X. Nuclear extracts were made as previously described^55^. Briefly, cells were lysed in buffer A (10mM HEPES, pH 7.9, 2.5 mM MgCl2, 0.25 M Sucrose, 0.1% NP-40 0.5 mM DTT, with protease inhibitors (Sigma #P8849) for 10 min at 4°C, and nuclei separated by centrifugation at 7,000xg for 10 min. Nuclei were washed twice with buffer A to reduce cytoplasmic and Cultrex contamination and resuspended in buffer B (25 mM HEPES, pH7.9, 1.5 M MgCl2, 700 mM NaCl, 0.5 mM DTT, 0.1 mM EDTA, 20% glycerol, protease inhibitor cocktail). After sonication, nuclear extracts were cleared by centrifugation and diluted to 2 ug/ul and 150mM NaCl. 500ug of extract were used for each IP. Extracts were pre-cleared with 25ul of magnetic streptavidin beads (Pierce #88816) for 1h at RT, followed by incubation with 2 ug of biotinylated nucleosomes overnight a 4°C. 25ul of magnetic streptavidin beads were then added to each IP and incubated for 1hr at RT. Using a magnetic stand, streptavidin beads washed once with IP buffer (20mM HEPES, pH 7.5, 150 mM NaCl, 5% Glycerol 1 mM EDTA) supplemented with 0.5% NP-40 and protease inhibitors, twice with 1x volume of RT protein wash buffer (20 mM HEPES, pH 7.5, 250 mM NaCl, 5% Glycerol 1mM DTT, 0.5% NP-40), and 3 times with 4°C MS wash buffer (20 mM NH3HCO3, pH8, 2 mM CaCl2). 1/10 of the sample was used for WB validation, the remaining beads were flash frozen in liquid nitrogen and stored at -80°C.

### V5 and H2A.Z IP-MS

Each IP was performed in triplicate. Nuclei were prepared from H9 human embryonic stem cells (hESCs) by lysing cells in Buffer A (10 mM HEPES pH 7.9, 2.5 mM MgCl2, 0.25 M sucrose, 0.1% NP40, 0.5 mM DTT, and protease inhibitors) and centrifugating 7000xg for 10 min. To remove cytoplasmic contamination, nuclei were washed 3 times in buffer A. After centrifugation, nuclei were resuspended in MNase lysis buffer (10 mM HEPES pH 7.9, 5 mM CaCl₂, 150 mM NaCl, plus EDTA/EGTA-free protease inhibitors; Sigma #P8849) and treated with MNase (Sigma-Aldrich, N3755) at 5 U per ∼10⁶ nuclei for 30 min at 37°C. Reactions were quenched with 1 mM EGTA, and lysates were supplemented with 0.5 mM DTT, protease inhibitors, and 0.01% NP-40. After full speed centrifugation, the supernatant was collected as the nuclear extract. MNase digestion was confirmed by DNA agarose gel after phenol-chloroform extraction. For V5 pulldowns, 25 µL of V5 nanobody conjugated beads (Proteintech #V5TMA) were incubated with 500 µg of nuclear extract for 1 h at 4°C for each reaction, except for H2A.Z.2–V5 samples, which were incubated with 5 mg of extract to compensate for lower expression. For H2A.Z IPs, 70 µL of Protein G Dynabeads (#10003) were incubated with 6 µL of H2A.Z antibody (Abcam #ab4174) for 30 min at 4 °C prior to addition to nuclear extracts. Beads were collected on a magnetic rack, washed first 3X with MNase lysis buffer containing 0.01% NP-40, then 2X protein wash buffer (20 mM HEPES pH 7.5, 300 mM NaCl, 5% glycerol, 0.5 mM DTT, 0.01% NP-40), and 2X ice-cold MS wash buffer (20 mM ammonium bicarbonate pH 8, 2 mM CaCl₂). 1/12 of the sample was used for WB validation, the remaining beads were flash frozen in liquid nitrogen and stored at -80C. A list of antibodies used is provided in Table S3.

### RNA-Seq

Total RNA was isolated from 1.5 × 10^6^ of H9 cells per biological replicate using a RNeasy Mini Kit (Quiagen #74104) according to the manufacturer’s protocol. PolyA enrichment, library preparation, and sequencing were performed by Novogene. The fastq files were aligned to the hg38 genome (GRCh38.p14) using STAR^56^. The sam files were converted to bam files using samtools^57^. The mapped reads were assigned to genes using the featureCounts function in Rsubread^58^. The count tables generated were used as an input for DESeq2 in R^59^ for differential gene expression analysis between the WT and mutant cells. Genes with a log2 fold change ≤-0.5 or ≥0.5 and FDR ≤ 0.05 were used for downstream analysis. Gene ontology analysis was performed using DAVID. PCA and volcano plot of the differentially expressed genes were made using ggplot2; other plots were made using GraphPad Prism.

### TMT-labeling and MS

Beads containing the immunoprecipitated samples were resuspended in digestion buffer (50 mM NH_3_HCO_3_, pH8), reduced with DTT (100 mM for 30 min at RT), alkylated with iodoacetamide (100 mM for 10 min at RT), and digested with two trypsin treatments at 37°C (500 ng overnight, followed by 500 ng for 4h). The supernatants were combined and desalted on C18 ZipTips and dried. Samples were labeled with either TMT-6plex reagents or TMTpro-18plex reagents depending on the experiment. TMT-6plex labeling was conducted by suspending the peptides in 10µl of 250 mM HEPES (pH 8.5) and adding 40µg reagent from 10 µg/µl stock in acetonitrile. Ater 1 hour of labeling, an additional 40µg aliquot of reagent was added and the labeling repeated. TMTpro-18plex labeling was conducted with a 96-well format kit. Each well contained 50µg reagent in DMSO and the labeling was performed according to the manufacturers protocol. Labeling efficiency was checked prior to final mixing and determined to be between 98.5-100%. Labeled peptides were then mixed in equal proportion, the organic component was removed or diluted and the samples were desalted and evaporated to dryness. Samples were brought up in 0.1% formic acid and mass spectrometry was performed on an Orbitrap Exploris 480 coupled through an EASY-Spray nano ion source to a Dionex UltiMate 3000 uPLC running an EASY-Spray column (75µm × 50cm column packed with 2µm, 100 Å PepMap C18 resin). Gradient and acquisition method details are as previously reported^60^.

Peak lists were generated using PAVA software^61^. The peaklists were searched against the human subset of the SwissProt database (SwissProt.2024.01.24), using Protein Prospector v 6.8.0^62^ with the following parameters: Enzyme specificity was tryptic with up to 2 missed cleavages per peptide. Carbamidomethylation of cysteine residues, and either TMT6plex or TMTPro16plex labeling of lysine residues and N terminus of the protein were allowed as fixed modifications. N-acetylation of the N terminus of the protein, loss of protein N-terminal methionine, pyroglutamate formation from of peptide N-terminal glutamines, and oxidation of methionine were allowed as variable modifications. Mass tolerance was 12 ppm for precursor ions and 25 ppm for product ions. The 50 most intense peaks were used for searching. The false discovery rate (FDR) was estimated by searching the data using a concatenated sequence database containing the original protein sequences plus randomized versions of each entry. Results were initially reported at a 1% FDR at the protein and peptide level but the FDR fell to essentially 0% after additional acceptance and filtering criteria were applied. Data preparation steps were performed in the R statistical computing environment. Initial protein inference reported all accession numbers that could be explained by the identified peptide sequences. The list was then filtered to a minimum set of accession numbers that would explain the set of peptides identified where only accession numbers identified by at least one unique peptide sequence, and at least two total peptide sequences were retained. If multiple accession numbers matched these criteria, one was selected at random.

For quantitation only unique peptides were considered; peptides common to several proteins were not used for quantitative analysis. Relative quantization of peptide abundance was performed via calculation of the intensity of reporter ions corresponding to the different TMT labels, present in product ion spectra. Intensities were determined by Protein Prospector. Protein level summarization, normalization, imputation, and statistical testing was performed using the MSstats and MSstatsTMT R packages^63^. Every IP condition and negative control was repeated with at least 3 biological replicates. For CRISPR edited hESC anti-H2AZ IPs, three monoclonal cell lines per condition were selected and three biological replicates from each clone were immunoprecipitated and TMTpro labeled. Enrichment analysis and plots were made using STRING^64^.

### V5 and H2A.Z CUT&Tag

CUT&Tag was performed with Active Motif assay kit (#53160) using *Drosophila* nuclei spike-in (#53168; #53173) following the manufacturer’s protocol. For each sample, 2.5×10^5^ H9 cells were mixed with 2.5×10^4^ *Drosophila* nuclei and incubated with anti-V5 (ThermoFisher, #R960-25) or anti-H2A.Z antibody (Active Motif #39013) together with the spike-in antibody. After tagmentation, libraries were PCR-amplified and sequenced on an Illumina NextSeq 2000. Reads were aligned to hg38 and dm6 genome references using Bowtie2, BAM files were generated using SAMtools and converted to bedgraph format using deepTools using a per-sample spike-in scale factor = [lowest Drosophila uniquely mapped reads across all sample/ number of uniquely mapped Drosophila reads in target sample]. Peaks were called using SEACR with the bedgraph files in stringent mode with the numeric threshold set to 0.05, skipping normalization. The significance of the signal difference between conditions was determined using bootstrap resampling.

### ATAC-seq

Cryopreserved cells were thawed in a 37°C water bath, pelleted, washed with cold PBS, counted and tagmented with a Tn5 enzyme. Briefly, cell pellets were resuspended in lysis buffer, pelleted, and tagmented using the enzyme and buffer provided in the ATAC-Seq Kit (Active Motif). Tagmented DNA was then purified using a DNA extraction method and then PCR amplified. The resultant libraries were purified using SPRI beads. The final libraries were quantified and sequenced with PE50 sequencing on the Illumina platform NovaSeq X and NextSeq 2K. Utilizing featureCounts, we obtained the count matrix where each row is a consensus peak region, and each column is a sample. We then perform differential analyses to identify statistically significant differential peaks using DESeq2^59^. A peak region is marked as differentially accessible if the BH-adjusted p value < 0.05 and |LFC| >1.

## References

1. McGinty, R.K., and Tan, S. (2015). Nucleosome Structure and Function. Chem. Rev. 115, 2255–2273. 10.1021/cr500373h.

2. Bannister, A.J., and Kouzarides, T. (2011). Regulation of chromatin by histone modifications. Cell Res. 21, 381–395. 10.1038/cr.2011.22.

3. Nakanishi, S., Sanderson, B.W., Delventhal, K.M., Bradford, W.D., Staehling-Hampton, K., and Shilatifard, A. (2008). A comprehensive library of histone mutants identifies nucleosomal residues required for H3K4 methylation. Nat. Struct. Mol. Biol. 15, 881–888. 10.1038/nsmb.1454.

4. Bagert, J.D., Mitchener, M.M., Patriotis, A.L., Dul, B.E., Wojcik, F., Nacev, B.A., Feng, L., Allis, C.D., and Muir, T.W. (2021). Oncohistone mutations enhance chromatin remodeling and alter cell fates. Nat. Chem. Biol. 17, 403–411. 10.1038/s41589-021-00738-1.

5. Nacev, B.A., Feng, L., Bagert, J.D., Lemiesz, A.E., Gao, J., Soshnev, A.A., Kundra, R., Schultz, N., Muir, T.W., and Allis, C.D. (2019). The expanding landscape of ‘oncohistone’ mutations in human cancers. Nature 567, 473–478. 10.1038/s41586-019-1038-1.

6. Bryant, L., Li, D., Cox, S.G., Marchione, D., Joiner, E.F., Wilson, K., Janssen, K., Lee, P., March, M.E., Nair, D., et al. (2020). Histone H3.3 beyond cancer: Germline mutations in Histone 3 Family 3A and 3B cause a previously unidentified neurodegenerative disorder in 46 patients. Sci. Adv. 6, eabc9207. 10.1126/sciadv.abc9207.

7. Khazaei, S., Chen, C.C.L., Andrade, A.F., Kabir, N., Azarafshar, P., Morcos, S.M., França, J.A., Lopes, M., Lund, P.J., Danieau, G., et al. (2023). Single substitution in H3.3G34 alters DNMT3A recruitment to cause progressive neurodegeneration. Cell 186, 1162–1178.e20. 10.1016/j.cell.2023.02.023.

8. Buschbeck, M., and Hake, S.B. (2017). Variants of core histones and their roles in cell fate decisions, development and cancer. Nat. Rev. Mol. Cell Biol. 18, 299–314. 10.1038/nrm.2016.166.

9. Wong, L.H., and Tremethick, D.J. (2025). Multifunctional histone variants in genome function. Nat. Rev. Genet. 26, 82–104. 10.1038/s41576-024-00759-1.

10. Maze, I., Noh, K.-M., Soshnev, A.A., and Allis, C.D. (2014). Every amino acid matters: essential contributions of histone variants to mammalian development and disease. Nat. Rev. Genet. 15, 259–271. 10.1038/nrg3673.

11. Dryhurst, D., Ishibashi, T., Rose, K.L., Eirín-López, J.M., McDonald, D., Silva-Moreno, B., Veldhoen, N., Helbing, C.C., Hendzel, M.J., Shabanowitz, J., et al. (2009). Characterization of the histone H2A.Z-1 and H2A.Z-2 isoforms in vertebrates. BMC Biol. 7, 86. 10.1186/1741-7007-7-86.

12. Greenberg, R.S., Long, H.K., Swigut, T., and Wysocka, J. (2019). Single Amino Acid Change Underlies Distinct Roles of H2A.Z Subtypes in Human Syndrome. Cell 178, 1421–1436.e24. 10.1016/j.cell.2019.08.002.

13. Horikoshi, N., Sato, K., Shimada, K., Arimura, Y., Osakabe, A., Tachiwana, H., Hayashi-Takanaka, Y., Iwasaki, W., Kagawa, W., Harata, M., et al. (2013). Structural polymorphism in the L1 loop regions of human H2A.Z.1 and H2A.Z.2. Acta Crystallogr. D Biol. Crystallogr. 69, 2431–2439. 10.1107/S090744491302252X.

14. Dunn, C.J., Sarkar, P., Bailey, E.R., Farris, S., Zhao, M., Ward, J.M., Dudek, S.M., and Saha, R.N. (2017). Histone Hypervariants H2A.Z.1 and H2A.Z.2 Play Independent and Context-Specific Roles in Neuronal Activity-Induced Transcription of Arc/Arg3.1 and Other Immediate Early Genes. eNeuro 4, ENEURO.0040-17.2017. 10.1523/ENEURO.0040-17.2017.

15. Giaimo, B.D., Ferrante, F., Herchenröther, A., Hake, S.B., and Borggrefe, T. (2019). The histone variant H2A.Z in gene regulation. Epigenetics Chromatin 12, 37. 10.1186/s13072-019-0274-9.

16. Guillemette, B., and Gaudreau, L. (2006). Reuniting the contrasting functions of H2A.Z. Biochem. Cell Biol. Biochim. Biol. Cell. 84, 528–535. 10.1139/o06-077.

17. Suto, R.K., Clarkson, M.J., Tremethick, D.J., and Luger, K. (2000). Crystal structure of a nucleosome core particle containing the variant histone H2A.Z. Nat. Struct. Biol. 7, 1121–1124. 10.1038/81971.

18. Hondele, M., Stuwe, T., Hassler, M., Halbach, F., Bowman, A., Zhang, E.T., Nijmeijer, B., Kotthoff, C., Rybin, V., Amlacher, S., et al. (2013). Structural basis of histone H2A–H2B recognition by the essential chaperone FACT. Nature 499, 111–114. 10.1038/nature12242.

19. D’Arcy, S., Martin, K.W., Panchenko, T., Chen, X., Bergeron, S., Stargell, L.A., Black, B.E., and Luger, K. (2013). Chaperone Nap1 Shields Histone Surfaces Used in a Nucleosome and Can Put H2A-H2B in an Unconventional Tetrameric Form. Mol. Cell 51, 662–677. 10.1016/j.molcel.2013.07.015.

20. Dias, J.K., Dias, P.S., Alakenova, R., Mariasoosai, C., Claridy, C., Gazi, S., Torabifard, H., and D’Arcy, S. (2026). Histone variant H2A.Z enhances histone and nucleosome dynamics. Mol. Cell. Proteomics, 101518. 10.1016/j.mcpro.2026.101518.

21. Iwasaki, W., Miya, Y., Horikoshi, N., Osakabe, A., Taguchi, H., Tachiwana, H., Shibata, T., Kagawa, W., and Kurumizaka, H. (2013). Contribution of histone N-terminal tails to the structure and stability of nucleosomes. FEBS Open Bio 3, 363–369. 10.1016/j.fob.2013.08.007.

22. Li, Z., and Kono, H. (2016). Distinct Roles of Histone H3 and H2A Tails in Nucleosome Stability. Sci. Rep. 6, 31437. 10.1038/srep31437.

23. Li, S., Wei, T., and Panchenko, A.R. (2023). Histone variant H2A.Z modulates nucleosome dynamics to promote DNA accessibility. Nat. Commun. 14, 769. 10.1038/s41467-023-36465-5.

24. Dryhurst, D., Ishibashi, T., Rose, K.L., Eirín-López, J.M., McDonald, D., Silva-Moreno, B., Veldhoen, N., Helbing, C.C., Hendzel, M.J., Shabanowitz, J., et al. (2009). Characterization of the histone H2A.Z-1 and H2A.Z-2 isoforms in vertebrates. BMC Biol. 7, 86. 10.1186/1741-7007-7-86.

25. Eirín-López, J.M., González-Romero, R., Dryhurst, D., Ishibashi, T., and Ausió, J. (2009). The evolutionary differentiation of two histone H2A.Z variants in chordates (H2A.Z-1 and H2A.Z-2) is mediated by a stepwise mutation process that affects three amino acid residues. BMC Evol. Biol. 9, 31. 10.1186/1471-2148-9-31.

26. Vardabasso, C., Gaspar-Maia, A., Hasson, D., Pünzeler, S., Valle-Garcia, D., Straub, T., Keilhauer, E.C., Strub, T., Dong, J., Panda, T., et al. (2015). Histone Variant H2A.Z.2 Mediates Proliferation and Drug Sensitivity of Malignant Melanoma. Mol. Cell 59, 75–88. 10.1016/j.molcel.2015.05.009.

27. Vardabasso, C., Hake, S.B., and Bernstein, E. (2016). Histone variant H2A.Z.2: A novel driver of melanoma progression. Mol. Cell. Oncol. 3, e1073417. 10.1080/23723556.2015.1073417.

28. Sanulli, S., Trnka, M.J., Dharmarajan, V., Tibble, R.W., Pascal, B.D., Burlingame, A.L., Griffin, P.R., Gross, J.D., and Narlikar, G.J. (2019). HP1 reshapes nucleosome core to promote phase separation of heterochromatin. Nature 575, 390–394. 10.1038/s41586-019-1669-2.

29. Gibson, B.A., Doolittle, L.K., Schneider, M.W.G., Jensen, L.E., Gamarra, N., Henry, L., Gerlich, D.W., Redding, S., and Rosen, M.K. (2019). Organization of Chromatin by Intrinsic and Regulated Phase Separation. Cell 179, 470–484.e21. 10.1016/j.cell.2019.08.037.

30. Ruhl, D.D., Jin, J., Cai, Y., Swanson, S., Florens, L., Washburn, M.P., Conaway, R.C., Conaway, J.W., and Chrivia, J.C. (2006). Purification of a Human SRCAP Complex That Remodels Chromatin by Incorporating the Histone Variant H2A.Z into Nucleosomes. Biochemistry 45, 5671–5677. 10.1021/bi060043d.

31. Mao, Z., Pan, L., Wang, W., Sun, J., Shan, S., Dong, Q., Liang, X., Dai, L., Ding, X., Chen, S., et al. (2014). Anp32e, a higher eukaryotic histone chaperone directs preferential recognition for H2A.Z. Cell Res. 24, 389–399. 10.1038/cr.2014.30.

32. Brahma, S., Udugama, M.I., Kim, J., Hada, A., Bhardwaj, S.K., Hailu, S.G., Lee, T.-H., and Bartholomew, B. (2017). INO80 exchanges H2A.Z for H2A by translocating on DNA proximal to histone dimers. Nat. Commun. 8, 15616. 10.1038/ncomms15616.

33. Lamaa, A., Humbert, J., Aguirrebengoa, M., Cheng, X., Nicolas, E., Côté, J., and Trouche, D. (2020). Integrated analysis of H2A.Z isoforms function reveals a complex interplay in gene regulation. eLife 9, e53375. 10.7554/eLife.53375.

34. Pünzeler, S., Link, S., Wagner, G., Keilhauer, E.C., Kronbeck, N., Spitzer, R.M., Leidescher, S., Markaki, Y., Mentele, E., Regnard, C., et al. (2017). Multivalent binding of PWWP2A to H2A.Z regulates mitosis and neural crest differentiation. EMBO J. 36, 2263–2279. 10.15252/embj.201695757.

35. Fujimoto, S., Seebart, C., Guastafierro, T., Prenni, J., Caiafa, P., and Zlatanova, J. (2012). Proteome analysis of protein partners to nucleosomes containing canonical H2A or the variant histones H2A.Z or H2A.X. Biol. Chem. 393, 47–61. 10.1515/BC-2011-216.

36. Mizuguchi, G., Shen, X., Landry, J., Wu, W.-H., Sen, S., and Wu, C. (2004). ATP-Driven Exchange of Histone H2AZ Variant Catalyzed by SWR1 Chromatin Remodeling Complex. Science 303, 343–348. 10.1126/science.1090701.

37. Creyghton, M.P., Markoulaki, S., Levine, S.S., Hanna, J., Lodato, M.A., Sha, K., Young, R.A., Jaenisch, R., and Boyer, L.A. (2008). H2AZ is enriched at polycomb complex target genes in ES cells and is necessary for lineage commitment. Cell 135, 649–661. 10.1016/j.cell.2008.09.056.

38. Calo, E., and Wysocka, J. (2013). Modification of Enhancer Chromatin: What, How, and Why? Mol. Cell 49, 825–837. 10.1016/j.molcel.2013.01.038.

39. Ku, M., Jaffe, J.D., Koche, R.P., Rheinbay, E., Endoh, M., Koseki, H., Carr, S.A., and Bernstein, B.E. (2012). H2A.Z landscapes and dual modifications in pluripotent and multipotent stem cells underlie complex genome regulatory functions. Genome Biol. 13, R85. 10.1186/gb-2012-13-10-r85.

40. Kreienbaum, C., Paasche, L.W., and Hake, S.B. (2022). H2A.Z’s ‘social’ network: functional partners of an enigmatic histone variant. Trends Biochem. Sci. 47, 909–920. 10.1016/j.tibs.2022.04.014.

41. Hu, G., Cui, K., Northrup, D., Liu, C., Wang, C., Tang, Q., Ge, K., Levens, D., Crane-Robinson, C., and Zhao, K. (2013). H2A.Z facilitates access of active and repressive complexes to chromatin in embryonic stem cell self-renewal and differentiation. Cell Stem Cell 12, 180–192. 10.1016/j.stem.2012.11.003.

42. Faast, R., Thonglairoam, V., Schulz, T.C., Beall, J., Wells, J.R.E., Taylor, H., Matthaei, K., Rathjen, P.D., Tremethick, D.J., and Lyons, I. (2001). Histone variant H2A.Z is required for early mammalian development. Curr. Biol. 11, 1183–1187. 10.1016/S0960-9822(01)00329-3.

43. Colino-Sanguino, Y., Clark, S.J., and Valdes-Mora, F. (2022). The H2A.Z-nucleosome code in mammals: emerging functions. Trends Genet. 38, 273–289. 10.1016/j.tig.2021.10.003.

44. Kazakevych, J., Sayols, S., Messner, B., Krienke, C., and Soshnikova, N. (2017). Dynamic changes in chromatin states during specification and differentiation of adult intestinal stem cells. Nucleic Acids Res. 45, 5770–5784. 10.1093/nar/gkx167.

45. Li, Z., Gadue, P., Chen, K., Jiao, Y., Tuteja, G., Schug, J., Li, W., and Kaestner, K.H. (2012). Foxa2 and H2A.Z Mediate Nucleosome Depletion during Embryonic Stem Cell Differentiation. Cell 151, 1608–1616. 10.1016/j.cell.2012.11.018.

46. Jin, C., Zang, C., Wei, G., Cui, K., Peng, W., Zhao, K., and Felsenfeld, G. (2009). H3.3/H2A.Z double variant–containing nucleosomes mark “nucleosome-free regions” of active promoters and other regulatory regions. Nat. Genet. 41, 941–945. 10.1038/ng.409.

47. Li, S., Wei, T., and Panchenko, A.R. (2023). Histone variant H2A.Z modulates nucleosome dynamics to promote DNA accessibility. Nat. Commun. 14, 769. 10.1038/s41467-023-36465-5.

48. Manglik, A., and Kruse, A.C. (2017). Structural Basis for G Protein-Coupled Receptor Activation. Biochemistry 56, 5628–5634. 10.1021/acs.biochem.7b00747.

49. Sinha, K.K., Gross, J.D., and Narlikar, G.J. (2017). Distortion of histone octamer core promotes nucleosome mobilization by a chromatin remodeler. Science 355, eaaa3761. 10.1126/science.aaa3761.

50. Bilokapic, S., Strauss, M., and Halic, M. (2018). Histone octamer rearranges to adapt to DNA unwrapping. Nat. Struct. Mol. Biol. 25, 101–108. 10.1038/s41594-017-0005-5.

51. Bilokapic, S., Strauss, M., and Halic, M. (2018). Structural rearrangements of the histone octamer translocate DNA. Nat. Commun. 9, 1330. 10.1038/s41467-018-03677-z.

52. Armache, J.P., Gamarra, N., Johnson, S.L., Leonard, J.D., Wu, S., Narlikar, G.J., and Cheng, Y. Cryo-EM structures of remodeler-nucleosome intermediates suggest allosteric control through the nucleosome. eLife 8, e46057. 10.7554/eLife.46057.

53. Dyer, P.N., Edayathumangalam, R.S., White, C.L., Bao, Y., Chakravarthy, S., Muthurajan, U.M., and Luger, K. (2004). Reconstitution of nucleosome core particles from recombinant histones and DNA. Methods Enzymol. 375, 23–44. 10.1016/s0076-6879(03)75002-2.

54. Costello, S.M., Shoemaker, S.R., Hobbs, H.T., Nguyen, A.W., Hsieh, C.-L., Maynard, J.A., McLellan, J.S., Pak, J.E., and Marqusee, S. (2022). The SARS-CoV-2 spike reversibly samples an open-trimer conformation exposing novel epitopes. Nat. Struct. Mol. Biol. 29, 229–238. 10.1038/s41594-022-00735-5.

55. Sanulli, S., Justin, N., Teissandier, A., Ancelin, K., Portoso, M., Caron, M., Michaud, A., Lombard, B., da Rocha, S.T., Offer, J., et al. (2015). Jarid2 Methylation via the PRC2 Complex Regulates H3K27me3 Deposition during Cell Differentiation. Mol. Cell 57, 769–783. 10.1016/j.molcel.2014.12.020.

56. Dobin, A., Davis, C.A., Schlesinger, F., Drenkow, J., Zaleski, C., Jha, S., Batut, P., Chaisson, M., and Gingeras, T.R. (2013). STAR: ultrafast universal RNA-seq aligner. Bioinforma. Oxf. Engl. 29, 15–21. 10.1093/bioinformatics/bts635.

57. Danecek, P., Bonfield, J.K., Liddle, J., Marshall, J., Ohan, V., Pollard, M.O., Whitwham, A., Keane, T., McCarthy, S.A., Davies, R.M., et al. (2021). Twelve years of SAMtools and BCFtools. GigaScience 10, giab008. 10.1093/gigascience/giab008.

58. Liao, Y., Smyth, G.K., and Shi, W. (2019). The R package Rsubread is easier, faster, cheaper and better for alignment and quantification of RNA sequencing reads. Nucleic Acids Res. 47, e47. 10.1093/nar/gkz114.

59. Love, M.I., Huber, W., and Anders, S. (2014). Moderated estimation of fold change and dispersion for RNA-seq data with DESeq2. Genome Biol. 15, 550. 10.1186/s13059-014-0550-8.

60. Smeyers, J., Oses-Prieto, J.A., Yadanar, L., Wang, M., Iadarola, M., Lu, S., Wang, K.S., Watanabe, T.H., Debnath, J., Burlingame, A.L., et al. (2025). Phospho-proteome profiling in human neurons reveals targets of TBK1 in ALS/FTD-associated autophagy networks. Cell Rep. 44, 116494. 10.1016/j.celrep.2025.116494.

61. Guan, S., Price, J.C., Prusiner, S.B., Ghaemmaghami, S., and Burlingame, A.L. (2011). A Data Processing Pipeline for Mammalian Proteome Dynamics Studies Using Stable Isotope Metabolic Labeling*. Mol. Cell. Proteomics 10, M111.010728. 10.1074/mcp.M111.010728.

62. Chalkley, R.J., Baker, P.R., Huang, L., Hansen, K.C., Allen, N.P., Rexach, M., and Burlingame, A.L. (2005). Comprehensive analysis of a multidimensional liquid chromatography mass spectrometry dataset acquired on a quadrupole selecting, quadrupole collision cell, time-of-flight mass spectrometer: II. New developments in Protein Prospector allow for reliable and comprehensive automatic analysis of large datasets. Mol. Cell. Proteomics MCP 4, 1194–1204. 10.1074/mcp.D500002-MCP200.

63. Figueroa-Navedo, A.M., Kapre, R., Gupta, T., Xu, Y., Phaneuf, C.G., Beltran, P.M.J., Xue, L., Ivanov, A.R., and Vitek, O. (2025). MSstatsTMT Improves Accuracy of Thermal Proteome Profiling. Mol. Cell. Proteomics 24. 10.1016/j.mcpro.2025.100999.

64. Szklarczyk, D., Kirsch, R., Koutrouli, M., Nastou, K., Mehryary, F., Hachilif, R., Gable, A.L., Fang, T., Doncheva, N.T., Pyysalo, S., et al. (2023). The STRING database in 2023: protein-protein association networks and functional enrichment analyses for any sequenced genome of interest. Nucleic Acids Res. 51, D638–D646. 10.1093/nar/gkac1000.

